# Size-dependent behavioral and antennal responses to doses of (+)-isopinocamphone and 1,8-cineole mixed with pheromone: a potential host selection strategy in female *Ips typographus* L.

**DOI:** 10.1101/2025.05.30.657003

**Authors:** Antonioni Acacio Campos Moliterno, Mayuri Kashinath Shewale, Sara Basile, Jiří Synek, Anna Jirošová

## Abstract

**Context:** *Ips typographus*, a major pest of Norway spruce (*Picea abies*) in Europe, is experiencing more frequent outbreaks due to climate change. These outbreaks involve shifts in population dynamics and phenotypic traits, influencing beetle responses to olfactory cues from stressed host trees.

**Aims:** The study examines the size-dependent behavioral and antennal responses of female *I. typographus* to two host selection–deciding volatiles with contrasting ecological roles: 1,8-cineole, which inhibits attraction to unsuitable trees, and (+)-isopinocamphone, a pheromone synergist. Size-linked morphological and olfactory adaptations may influence females’ ability to select suitable host trees for reproduction.

**Methods:** In field trap experiments conducted in 2019 and 2022, the body size of *I. typographus* females caught in response to different doses of (+)-isopinocamphone or 1,8-cineole in combination with pheromone was compared. Female *Ips typographus* were sorted based on body length, the size of the antennal club was measured, and size-dependent antennal responses to these volatiles were analyzed using electroantennography.

**Results:** Larger females were more attracted to (+)-isopinocamphone in combination with pheromone in the field, showed stronger antennal detection, and had proportionally larger antennal clubs than smaller females. In contrast, smaller females were less repelled by 1,8-cineole added to pheromone but, in contradiction, antennally detected it more strongly than larger females despite having smaller antennal clubs.

**Conclusion:** The total body length significantly influences semiochemical detection in *I. typographus* females. (+)-isopinocamphone was detected more effectively by larger females, implying an advantage in the selection of suitable host trees. In contrast, the discrepancy between behavioral and antennal responses to 1,8-cineole in smaller females suggests the involvement of not only peripheral detection but also central nervous processing of olfactory signals driving behavior. This adaptation may enable smaller females to reduce competition with large ones by selecting less suitable trees. These findings provide new insights into the ecological relationship between beetle morphology and olfactory cues, with implications for tree–bark beetle interactions.

**Key message:** This study revealed differing behavioral and antennal responses between large and small female *I. typographus* to two bioactive oxygenated monoterpenes, (+)-isopinocamphone and 1,8-cineole, which serve contrasting ecological roles as aggregation pheromone synergist and inhibitor. Larger females were more attracted to (+)-isopinocamphone and had larger antennal clubs leading to enhanced antennal sensitivity, potentially improving their ability to select suitable host trees. In contrast, smaller females were less repelled by 1,8-cineole but had higher antennal sensitivity despite having smaller antennae. This discrepancy can be explained by behavioral decisions made after downstream olfactory signal processing in the central nervous system (CNS) and by the co-localization of 1,8-cineole with pheromone-sensitive neurons. Ecologically, small females may avoid competition with larger females by selecting less suitable trees. In conclusion, females’ body size influences olfactory-driven response to potential host selection decisive volatiles, which can impact reproductive success and bark beetle population dynamics.

## 1. Introduction

The Eurasian spruce bark beetle, *Ips typographus* L. 1758 (Coleoptera: Curculionidae), is a major pest associated with the Norway spruce (*Picea abies*) in Europe (Hlásny et al. 2021; Powell et al. 2021). Outbreaks of this species have intensified in frequency and severity, mainly due to climate change, and are facilitated by its complex and sophisticated chemical communication system (Biedermann et al. 2019). Male *I. typographus* play a pivotal role in locating weakened or stressed spruce trees using a combination of visual and chemical cues across both long and short distances (Birgersson and Bergström 1989; Netherer et al. 2021; Lehmanski et al. 2023). After initiating attack by boring into the bark (Wermelinger 2004), males produce aggregation pheromones (Birgersson et al. 1984; Ramakrishnan et al. 2022) to attract conspecifics for coordinating mass attacks to colonize the host tree and overcome tree defenses (Franceschi et al. 2005; Raffa et al. 2016). The success of this colonization process is highly influenced by host-emitted volatile organic compounds.

Norway spruce releases a range of monoterpenes, including highly abundant compounds such as α-pinene, β-pinene, β-phellandrene, and limonene (Netherer et al. 2021). These compounds have been tested to enhance the attraction of *I. typographus* (Erbilgin et al. 2007; Hulcr et al. 2006). In addition to these dominant volatiles, many studies have identified several low-abundance compounds emitted by spruce, comprising approximately 1% of the total volatile profile. While present in low amounts, these compounds elicit strong antennal responses in beetles (Kalinová et al. 2014; Schiebe et al. 2019), highlighting their ecological significance in tri-trophic interactions with beetles, its symbiotic microbiota, and the host tree (Netherer et al. 2021). Most of these minor yet biologically active volatiles are oxygenated spruce monoterpenes, with a few exceptions such as estragole and styrene, which have phenolic character. Oxygenated monoterpenes are produced within the spruce–bark beetle–symbiotic microorganism niche through multiple mechanisms. They can be formed via the oxidation of major spruce monoterpene hydrocarbons, either naturally by air exposure or enzymatically by the spruce microbiome. This transformation becomes especially prominent when trees experience stress, such as after being cut, windthrown, or infested by bark beetles (Netherer at al.2021; Schiebe et al. 2019). Under these conditions, the levels of compounds such as isopinocamphone, camphor, pinocarvone, terpinen-4-ol, and terpineol significantly increase. However, they remain minor components of spruce volatile profile compared to the main terpenic hydrocarbons.

Additionally, monoterpene hydrocarbons can be hydroxylated (introducing oxygen to molecule) by the beetles’ enzymatic detoxification systems. For example, α-pinene can be converted into myrtenol or *cis*-verbenol (Blomquist and Vogt 2021), while limonene may be transformed into carvone (Duetz et al. 2001). Over evolutionary time, several of these oxidation products have been co-opted by bark beetles as pheromonal compounds e.g. *cis*-verbenol in *I. typographus* (Francke and Vite 1983). Moreover, the beetle-associated intestinal microbiome also plays a key role in modifying host volatiles. It contributes not only to the oxidation of tree hydrocarbons but also to the further oxidation of *cis*-verbenol into verbenone, a potent bark beetle anti-aggregation signal (Frühbort et al. 2023). In parallel, beetle-exosymbiotic ophiostomatoid fungi, which are inoculated into trees by boring beetles during colonization, also metabolize monoterpenes to their oxidative forms (Kandasamy et al. 2023). On the other hand, some oxygenated monoterpenes, namely 1,8-cineole, are directly *de novo* formed through the cyclization of oxygenated intermediates within the spruce tree enzymatic system and not by oxidation of hydrocarbon precursors (Celedon and Bohlmann 2019).

Like many insects, bark beetles depend on highly specialized olfactory systems located in their antennae to navigate and interact with their environment (Hansson and Stensmyr 2011). In *I. typographus*, olfactory sensory neurons (OSNs) are housed within hair-like sensilla on the antennal surface. These neurons enable precise discrimination among a wide array of odor cues, including aggregation pheromones, host- and nonhost-derived tree volatiles, and volatiles produced by symbiotic microorganisms (Andersson et al. 2009). OSNs differ in their specificity. Some range from highly selective specialists detecting specific pheromones (Wojtasek et al. 1998) while some are broadly tuned generalists responsive to diverse environmental cues like host volatiles (Andersson et al. 2010; Binyameen et al. 2014). This specificity is determined by odorant receptors (ORs) located on their dendrites (Carey et al. 2010; Hallem and Carlson 2006). Upon odor detection, signals are transmitted to the antennal lobes (ALs), where glomeruli integrate input (Vosshall et al. 2000), and projection neurons relay this information to the mushroom bodies, which are involved in learning and memory, and the lateral horn, associated with innate behavioral responses (Galizia 2014; Clark and Ray 2016). This finely tuned chemosensory system plays a critical role in mediating behaviors such as host location, mate finding, and avoidance of unsuitable environments (Andersson et al. 2009; Zhang and Schlyter 2004).

Functional mapping of these neurons in *I. typographus* has identified specific OSN classes that respond to oxygenated spruce monoterpenes, including 1,8-cineole and (+)-isopinocamphone ((+)-IPC) (Andersson 2012; Kandasamy et al. 2023). Interestingly, OSNs activated by 1,8-cineole are consistently co-localized within the same sensilla as those tuned to the pheromonal component *cis*-verbenol (Andersson et al. 2009; Andersson et al. 2010). This arrangement of OSNs enables peripheral-level signal integration, where exposure to high concentrations of 1,8-cineole suppresses the neural response to *cis*-verbenol (Andersson et al. 2009; Binyameen et al. 2014). In contrast, OSNs responsive to (+)-isopinocamphone in *I. typographus* are individually localized and have not been observed in co-localization with neurons detecting other compounds. The specificity of this response is attributed to the olfactory receptor ItypOR29, located on the OSN membrane, which binds selectively to (+)-isopinocamphone, as confirmed through receptor expression and functional characterization in *I. typographus* (Hou et al. 2021).

Behavioral studies further support the ecological relevance of these olfactory interactions of oxygenated spruce monoterpenes. Field experiments have shown that 1,8-cineole, when added to pheromone blends containing *cis*-verbenol, inhibits beetle attraction in a clear dose-dependent manner (Andersson et al. 2010; Jirošová et al. 2022). Moreover, studies on the functional role of neuronal co-localization, where one neuron within the same sensillum responds to an attractant and another to an inhibitor, have demonstrated that 1,8-cineole induces more precise spatial avoidance of beetles from the pheromone source than verbenone. Verbenone is a known anti-attractant (Frühbort et al. 2023), yet its corresponding neuron has never been found to be co-localized with those for pheromonal compounds (Binyameen et al. 2014). Interestingly, a significantly higher content of 1,8-cineole has been found in spruce trees that are less susceptible to bark beetle attacks or that survived infestations more successfully (Schiebe et al. 2012). Additionally, preliminary feeding studies further indicate that 1,8-cineole exhibits greater toxicity to female *I. typographus* than to males (Zaman et al. 2024). This suggests that 1,8-cineole could serve as a potential chemical marker of bark beetle-resistant trees. In contrast to the inhibitory effects of 1,8-cineole, field studies showed that (+)- isopinocamphone significantly enhanced *I. typographus* captures at pheromone-baited traps and acted as a synergist with pheromone activity (Moliterno et al. 2023). Among the several tested compounds including estragole, 1,8-cineole, (±)-camphor, (−)-carvone, α-terpineol, (−)- terpinen-4-ol, (+)-pinocamphone, and (+)-isopinocamphone, each evaluated at three different doses; (+)-isopinocamphone was the only one to exhibit this relatively rare synergistic effect with pheromone. Additionally, (+)-isopinocamphone was also identified as a substantial component of the volatile bouquet produced by *Grossmania penicillata, Leptographium europhioides* and *Ophiostoma bicolor* which are beetle-associated fungi, when cultured on spruce phloem media. This fungal volatile blend was shown to attract beetles in short-range Petri-dish bioassays (Kandasamy et al. 2023).

Variations in abiotic and biotic factors significantly influence tree physiology, which directly affects host suitability and selection by bark beetles (Netherer et al. 2024). During endemic population stages, beetles prefer high-quality trees with low competition, conditions that favor offspring growth and fitness. However, during epidemic outbreaks, beetles are often forced to colonize suboptimal hosts, leading to reduced offspring vigor, including smaller body size (Foelker and Hofstetter 2014; Sallé and Raffa 2007). This decline in body size has cascading effects, as it can negatively influence pheromone production (Anderbrant et al. 1985; Pureswaran and Borden 2003), ultimately reducing mating success (Dacquin et al. 2024). The reproductive biology of *I. typographus* is closely linked to chemical signals. Males produce pheromones that serve not only for aggregation but also function partially as sexual attractants. As polygynous species, males typically mate with up to four females, increasing their mating success and overall fecundity (Schebeck et al. 2023). Female beetles are central to reproductive success, as they are responsible for gallery construction and oviposition. Consequently, females play a more selective role in reproduction, evaluating both mate quality and host tree suitability to optimize offspring survival and fitness (Schlyter and Zhang 1996, Paynter et al. 1990), relying on signals from both male-produced pheromones and tree-emitted volatile cues. The precision of pheromone-based recognition is well documented at the interspecific level among *Ips* beetles, suggesting strong selective pressures on olfactory systems (Schlyter et al. 2015). Variations in female total body length, combined with pheromonal and host tree chemical cues, are crucial for understanding ecological adaptations, such as host selection strategies (Muller et al. 2020; Schlyter and Anderbrant 1993).

This study explores size-specific behavior in female *I. typographus*, with a focus on their olfactory assessment of host tree quality, which is a critical factor for survival and reproductive success. We examine whether large and small females respond differently to two oxygenated spruce monoterpenes, 1,8-cineole and (+)-isopinocamphone, tested in combination with aggregation pheromones in a trap-capturing experiment. Additionally, we investigate if their antennae exhibit size-dependent differences in sensitivity to these compounds using electroantennographic analysis, and whether the antennal club shape differs between large and small females. Building on these objectives and the current understanding of bark beetle chemical ecology, this study is guided by the core hypothesis: "Larger and smaller *I. typographus* females will exhibit distinct behavioural and electrophysiological responses to the two oxygenated spruce monoterpenes with contrasting ecological roles: 1,8-cineole, which inhibits attraction to unsuitable trees, and (+)-isopinocamphone, which enhances the attraction to aggregation pheromones. These behavioral differences could be caused by size-dependent variations in antennal sensitivity for these two compounds, which is potentially influenced by morphological differences in the antennal clubs."

The ecological impact of these size-dependent differences, driven by morphological and olfactory adaptations, may affect *I. typographus* females’ ability to select high-quality host trees. This, in turn, could have broader implications for the beetles’ reproductive strategies and, consequently, their population dynamics.

## 2. Material and methods

### 2.1. Experimental approach

We evaluated the responses of *I. typographus* females using two complementary assays. The first was a field assay involving traps baited with either 1,8-cineole or (+)-isopinocamphone at three different doses in combination with a pheromone, and we compared the sex-ratio and body length of beetle captures to those from traps baited with pheromone alone. The second assay included electroantennography (EAG) analysis to measure the antennal responses of small and large females to varying doses of 1,8-cineole or (+)-isopinocamphone. We also conducted a morphometric analysis of antennal club size in these two groups.

### 2.2. Field experiment area and pheromone traps

The trapping experiments were conducted in 2019 and 2022 at the Forest CZU property in Kostelec nad Černými lesy, Czech Republic. The experiments took place in a mature, 100-year-old Norway spruce forest, a natural habitat for *I. typographus*, located at 600 meters above sea level. In 2019, the experiment was conducted at coordinates (49°56′02″N, 14°52′21″E), while the 2022 experiment took place at (49°55′57″N, 14°55′13″E). Both experiments were conducted during the same time frame: June 3 to July 28 of each year. In both the 2019 and 2021 experiments, traps were set up approximately 30 meters from the forest edge in a two-year-old clearing. They were arranged in a row, with a minimum distance of 15 meters between each trap, and were installed on wooden poles 1.5 meters above the ground.

In 2019, seven cross-vane Ecotraps (Fytofarm, Slovak Republic) were used for the collecting data for this experiment: three traps were baited with three different doses (low, medium, high) of 1,8-cineole or (+)-isopinocamphone, respectively, in combination with pheromone. One trap was baited with pheromone alone and served as a control (for baits composition see Table 1 for details). To minimize positional bias, the positions of the tested baits among these seven traps were changed seven times according to a randomization scheme based on a Latin square design (Evans et al. 2020).

**Table 1.**
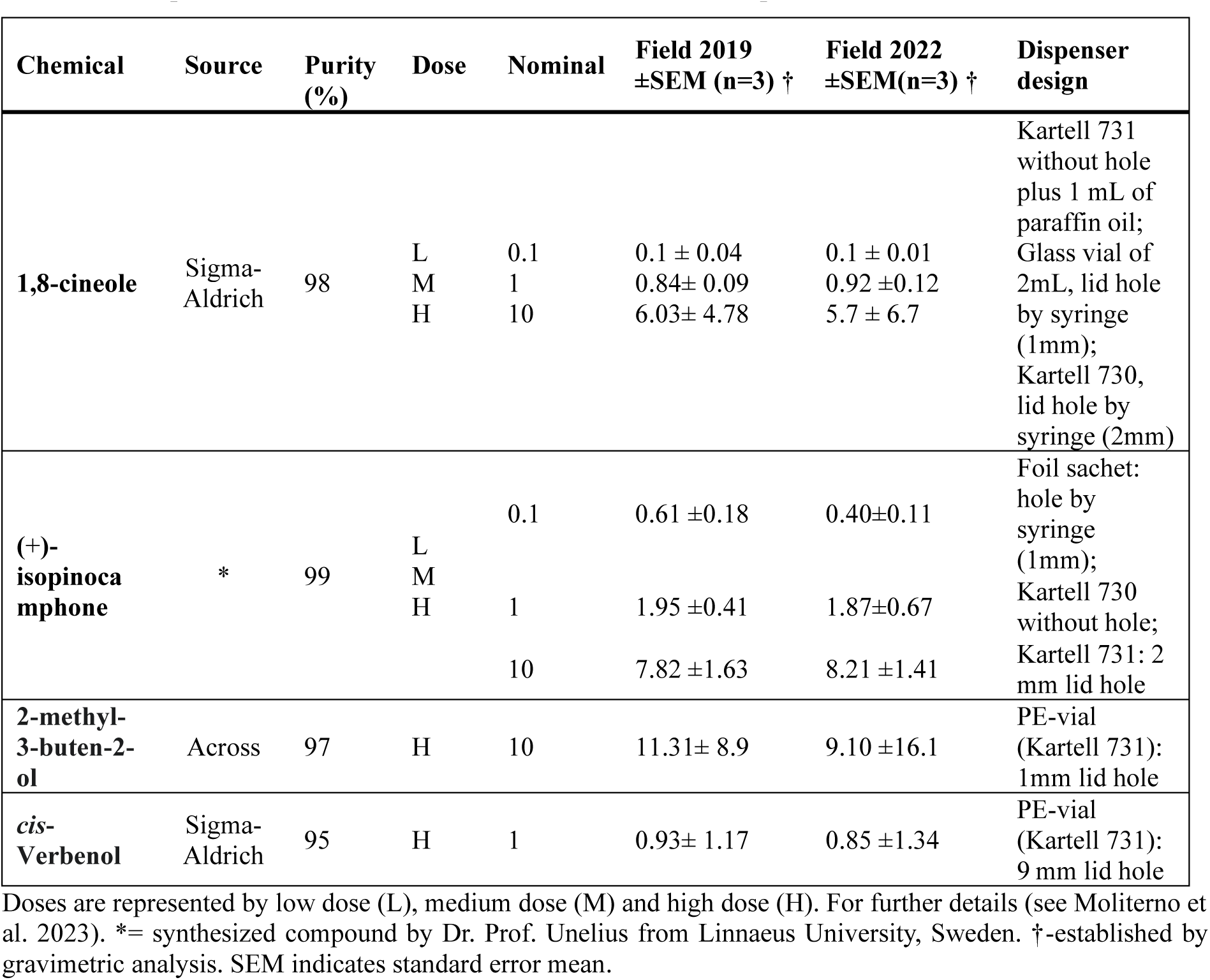
Description of treatment bait characteristics used in the experiments conducted in 2019 and 2022.

In 2022, for each compound (1,8-cineole and (+)-isopinocamphone), one block was set up, consisting of four traps arranged in a row: three traps baited with different doses of the tested compounds in combination with pheromone, and one trap with pheromone only (control). The positions of the tested baits among these four traps were changed four times according to a randomization scheme based on a Latin square design (Evans et al. 2020). These four rotations were repeated twice for each compound, resulting in a total of eight collections of catches for each treatment (Moliterno et al. 2023). Insects collected during the field experiment in both localities were preserved in ethanol for further analysis, including counting, sex sorting, and measurement.

### 2.3. Source and selection of beetles used for body length and antennal size measurement and electroantennographic detection analysis

For further measurement, fifty beetles were randomly selected from the ethanol-stored beetles caught in one of three doses of 1,8-cineole or (+)-isopinocamphone combined with pheromone, or caught with pheromone alone (a total of 8 groups each consisting of 50 randomly selected beetles). These beetles were selected from each replication of the experiments conducted in 2019 and 2021. Selected beetles were dried on tissue paper at 25°C for two hours, sorted by sex and measured for body length. Damaged specimens were excluded from the analysis.

For antennal club size measurements and electroantennography studies, *I. typographus* (F0 generation) emerged from naturally infested Norway spruce logs (*n*= 12; ±50 x 28 cm) collected in Kostelec nad Černými Lesy from June to July 2024 were used. The beetles were collected by placing naturally infested Norway spruce logs into fine mesh emergence cages under controlled laboratory conditions. The logs were monitored daily, and newly emerged adult beetles were collected manually from the mesh enclosures. Only females, ±3 days old, were selected after sorting by sex for further measurements and experiments.

### 2.4. Morphometric Analysis

The total body length of adult female *I. typographus* collected from field traps was measured in millimetres as demonstrated from the apical margin of the pronotum to the distal end of the elytra, using traditional linear morphometric analysis. The body size of the captured females was measured using a graticule (1–10 mm) integrated into a Nikon SMZ800N stereomicroscope at 30X magnification. Measurements were taken by the same researcher to ensure consistency, with recorded sizes ranging ranged from 4.2 to 5.3 mm millimetres. Based on this size range, two individuals were classified into two distinct size categories were established for further analysis. Female specimens selected for antennal club measurements and electroantennography were divided into:

1. Large-sized females: Body length ≥ 4.80 mm (n = 30)
2. Small-sized females: Body length ≤ 4.70 mm (n = 30)

To measure the antennal club measurements and electroantennography, excised antennae were mounted on borosilicate glass and imaged using a Nikon DFK 33UX250 camera (Imaging Source®, Germany) attached to a Nikon SMZ800N stereomicroscope. The antennal club length was measured from the apical end (ventral side) to the tip of the last antennomere, while the width was measured at the midpoint of the antennal club (ventral side). Measurements were obtained using IC Capture - Image Acquisition 4.0 software. The average measurements, calculated from the left and right antennae of each individual, were recorded in micrometres.

### 2.5. Electroantennographic (EAG) analysis

The sources and purity of the chemicals used for electroantennography (EAG) experiments were the same as described in Table 1. Dose-response tests were conducted using an aggregation pheromone in a 10:1 ratio of 2-methyl-3-buten-2-ol (MB) to *cis*-verbenol (cV), as well as the individual compounds 1,8-cineole and (+)-isopinocamphone. Antennae from large and small females were stimulated with odor stimuli at seven doses: 0.001 µg, 0.01 µg, 0.1 µg, 1 µg, 10 µg, 100 µg, and 1000 µg. For odor cartridge preparation, 10 µl of each odor stimuli solution at the corresponding concentration (diluted in hexane) was applied to a 1×1 cm strip of Whatman No. 1 filter paper. The solvent was allowed to evaporate for 1 minute before the strip was inserted into a glass Pasteur pipette (10 cm in length, 6 mm outer diameter), which was then used as an odor delivery cartridge for stimulation. Electrophysiological analyses were conducted using *I. typographus* females (F0 generation), as previously described. The F0 generation was chosen to directly represent the wild population of beetles originating from natural spruce forests. Prior to Analysis, insects were immobilized by cooling at 4°C for 5 minutes. This approach ensured the selection of morphometrically classified females within two size categories: large (≥ 4.80 mm, *n* = 10) and small (≤ 4.70 mm, *n* = 10).

Electroantennogram (EAG) analysis were conducted as described (Zhang et al. 2000). The sex of the beetles was determined through dissection, and the heads of female beetles were excised using a microblade. Two capillary glass electrodes filled with Ringer’s solution were used: one electrode was connected to the antennal club, while the other served as a reference by being inserted into the excised beetle head. The electrodes were attached to holders with an EAG probe (Syntech, Germany) and connected to a pre-amplifier. A constant stream of humidified air (200 ml/min) was directed over the antenna using a Syntech stimulus controller.

Odor cartridges (prepared as described above) were used to stimulate the antenna, and responses were recorded using EAG Pro software (Syntech, IDAC-4). Each stimulus (odor or control) was delivered as a 0.5-second pulse into the airstream directed at the antennal preparation, ensuring brief and consistent exposure. Control and odor stimuli were presented sequentially, with a one-minute interval between stimulations, allowing for antennal recovery and avoiding adaptation. The EAG probe was configured with a 0–32 Hz filter and a sampling rate of 100 Hz. Antennal responses were recorded as downward deflection signals in millivolts (mV), with response amplitudes defined as the peak depolarization of the olfactory sensilla of antennae measured during the 0.5-second odor stimulation. For each female beetle, recordings were made starting with the control stimulus and followed by sequential doses of the respective compound, increasing from the lowest to the highest concentration (0.001 µg to 1000 µg) to minimize sensory adaptation. Each biological replicate consisted of a single female beetle tested once per stimulus (*n* = 10 individuals per tested compound). The mean peak response amplitude across all replicates was calculated to assess antennal sensitivity to each compound.

### 2.6. Statistical Analysis

We tested normality within each treatment group from 2019 or 2022 using the Shapiro-Wilk test, and homogeneity of variances was assessed using Levene’s test. The raw data (total body length of adult female *I. typographus*) were exponentially transformed (Manly 1976), adjusting the assumption toward normality and equal variance. One-way ANOVAs were conducted separately for each year to assess whether insect body size differed significantly among treatment groups. Each ANOVA was followed by Tukey’s HSD test for post hoc comparisons, controlling the experiment-wise error rate. The Pearson’s Chi-squared test with Yates’ continuity correction was applied to check whether female proportion diverge regardless the dosage tested (Zar 2014).

The data obtained from small and large female *I. typographus* were compared using the Wilcoxon signed rank test (Harris and Hardin 2013; Hothorn et al. 2022). The two-sample analysis assessed differences in:

1. total body length between large and small females;
2. antennal club length between large and small females;
3. antennal club width between large and small females;

The chosen test deal with non-normality assumption as described before, but also paired-samples (the emerged adults insects collected from the same Norway spruce logs) and repeated measurement from antennal club length and width. The posterior analysis evaluated whether antennal club growth follows an isometric or allometric pattern relative to body size, we employed the Standardized Major Axis (SMA) regression using the "smatr" package in R (Warton, 2012). Before conducting an SMA regression, the length and width were log-transformed (ln= log natural), providing comparability, addressing potential scale issues, and making the relationship linear for better interpretation (Legendre and Legendre 1998). SMA regression was selected because it accounts for measurement errors in body size and antennal club length. SMA evaluates slopes >1 indicated isometric relationships, while deviations from <1 indicated allometry (Jolicoeur 1990; Warton et al. 2006). To evaluate the dose-response in electroantennography (EAG) analysis between large (*n* = 10) and small females (*n* = 10), the Wilcoxon signed rank test for repeated measurements was applied. All statistical analyses were performed using RStudio version 4.1.1 (Core R Team 2015), with a significance level (alpha) of 0.05. The dataset and R script used for the analysis are publicly available in the Dryad Digital Repository: https://doi.org/10.5061/dryad.rxwdbrvn1 (Moliterno et al. 2025). All figures were created using GraphPad Prism (version 9.5.0) software for macOS.

## 3. Results

We analyzed the sex ratio of beetles caught in the field using pheromone traps baited with three doses of 1,8-cineole and (+)-isopinocamphone in combination with pheromone, collected in 2019 (N=7) and 2022 (N=8) (Moliterno et al. 2023). In both years, females comprised 70–85% of the captures across treatments and pheromone-only groups (supplementary material, figure 1A and table 1A). In 2022, the proportion of females was significantly higher for all three doses of (+)-isopinocamphone combined with pheromone compared to the appropriate doses of 1,8-cineole combined with pheromone (Table 2, refer supplementary Table 1E and 1F for more details).

**Table 2.** Pearson’s Chi-squared test with Yates’ continuity correction comparing male and female *Ips typographus* catches for two compounds, (+)-isopinocamphone ((+)-IPC) and 1,8-cineole across different dose levels and years.

| Group | Absolute catches per females and compound |  | (+) -IPC vs 1,8-cineole |  |  |
| --- | --- | --- | --- | --- | --- |
|  | (+) -IPC females/total catches of beetles | 1,8-Cineole females/total catches of beetles | Chi-sq | df | p-value |
| 2019 - Low | 1267/1508 | 892/1059 | 0.008 | 1 | 0.9285 |
| 2019 - Medium | 2329/2875 | 1682/2103 | 0.755 | 1 | 0.3848 |
| 2019 - High | 2031/2477 | 294/370 | 1.217 | 1 | 0.2699 |
| 2022 - Low | 1052/1267 | 482/651 | 21.152 | 1 | <0.001** |
| 2022 - Medium | 1125/1406 | 389/519 | 5.488 | 1 | 0.0191* |
| 2022 - High | 1495/1917 | 183/258 | 6.028 | 1 | 0.0141* |

Data represents absolute beetle catches pooled from the respective number of trap rotations per year (2019: 7 rotations; 2022: 8 rotations). Chi-squared values indicate the results of contingency tests comparing female catches across treatments. Df represents degrees of freedom. p-values indicate the significance level of the observed differences between male respectively female catch proportions for respective compound and dose, with significance considered at *p* < 0.05 (*), *p* < 0.001 (**). Refer supplementary Table 1G for more details.

### 3.1. Prevalence and total body length differences in female captures across treatments and years

Females captured in control traps containing only pheromones had an average body length of 4.82 mm (SD = 0.22) in 2019 and 4.90 mm (SD = 0.17) in 2022. In contrast, females captured in traps baited with a high dose of 1,8-cineole were smaller, with an average body length of 4.69 mm (SD = 0.26), (*F* = 3.15, p = 0.026) in 2019 and 4.69 mm (SD = 0.23), (*F* = 9.59, p < 0.001) in 2022 (Fig. 1A). For (+)-isopinocamphone, trap catches showed significant differences based on dose. In 2019, females captured in traps baited with a low dose of (+)- isopinocamphone had an average body length of 4.75 mm (SD = 0.24) (*F* = 3.03, p = 0.03), while in 2022, the average was 4.75 mm (SD = 0.23) (*F* = 2.95, p = 0.03). Conversely, traps baited with a high dose of (+)-isopinocamphone attracted larger females, with average body lengths of 4.88 mm (SD = 0.19) in 2019 and 4.86 mm (SD = 0.20) in 2022 (Fig. 1B). Detailed data and statistical analyses, including results from ANOVA followed by Tukey’s HSD test (supplementary material, Table 1B) and visualized in Fig. 1.

**Figure 1.**
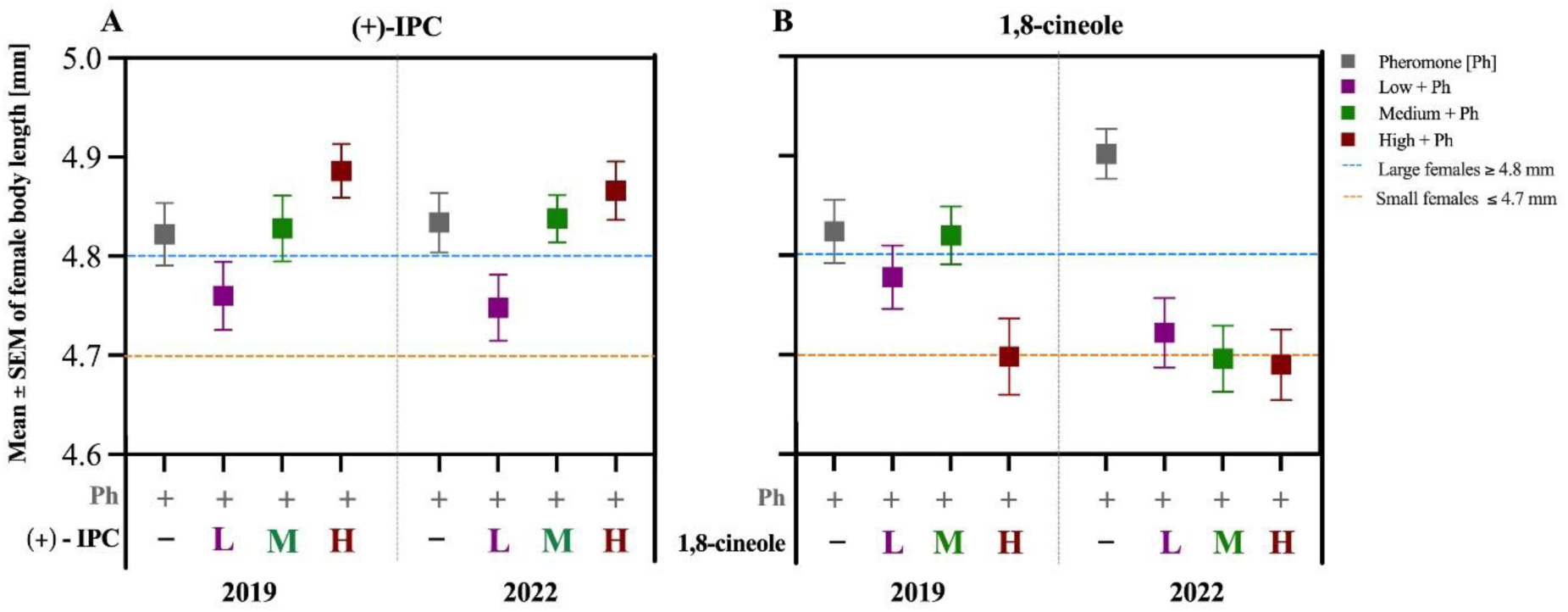
Mean body size of female *I. typographus* captured in response to three doses (low, medium, high) of (A) 1,8-cineole and (B) (+)-IPC is (+)-isopinocamphone, along with a control (pheromone only = Ph), in 2019 and 2022. Colors represent different doses or the control, with a sample size of *n* = 50 per group. Vertical bars show the standard error of the mean, and (*) indicates significant differences between groups based on Tukey’s HSD test (p = 0.05).

### 3.2. Total body length, antennal club size of large and small females is isometric to body size

The Wilcoxon signed rank for dependent sample analysis showed total body size (V= 465, p< 0.001), length, antennal club in length (V= 416, p< 0.001), and width (V= 431, p< 0.001) measurements differed significantly between large versus small females (Table 3).

**Table 3.** Means of measurements in total body length and antennal club (length and width) of large and small females of *Ips typographus;* mean ± SD.

| Parameters (mean ± SD) | Large females (N=30) | Small females (N=30) |
| --- | --- | --- |
| Body length (mm) | 4.88 ± 0.079 | 4.59 ± 0.105 |
| Antennal club length ($\mu\text{m}$) | 258.72 ± 20.05 | 233.59 ± 17.35 |
| Antennal club width ($\mu\text{m}$) | 222.76 ± 16.11 | 198.75 ± 14.94 |

Total body length data from females of *Ips typographus* was defined as large $\geq 4.80$ mm ( $n = 30$ ) versus small $\leq 4.70$ mm ( $n = 30$ ) and its respective antennal club measurements.

The standardized major axis (SMA) regression focusing on the correlation between length and width indicated significant and positive correlations (Large= R²= 0.43, p ≤ 0.001) and (Small= R²= 0.32, p= 0.001). The relationship between antennal club length and width log-transformed indicated that both were scaled isometrically, with slopes close to 1 (Large: 0.99, Small: 1.0) (supplementary material, Table 1C). This suggests a proportional relationship between length and width in both groups, where the two variables increase at similar rates (Fig. 2).

**Figure 2.**
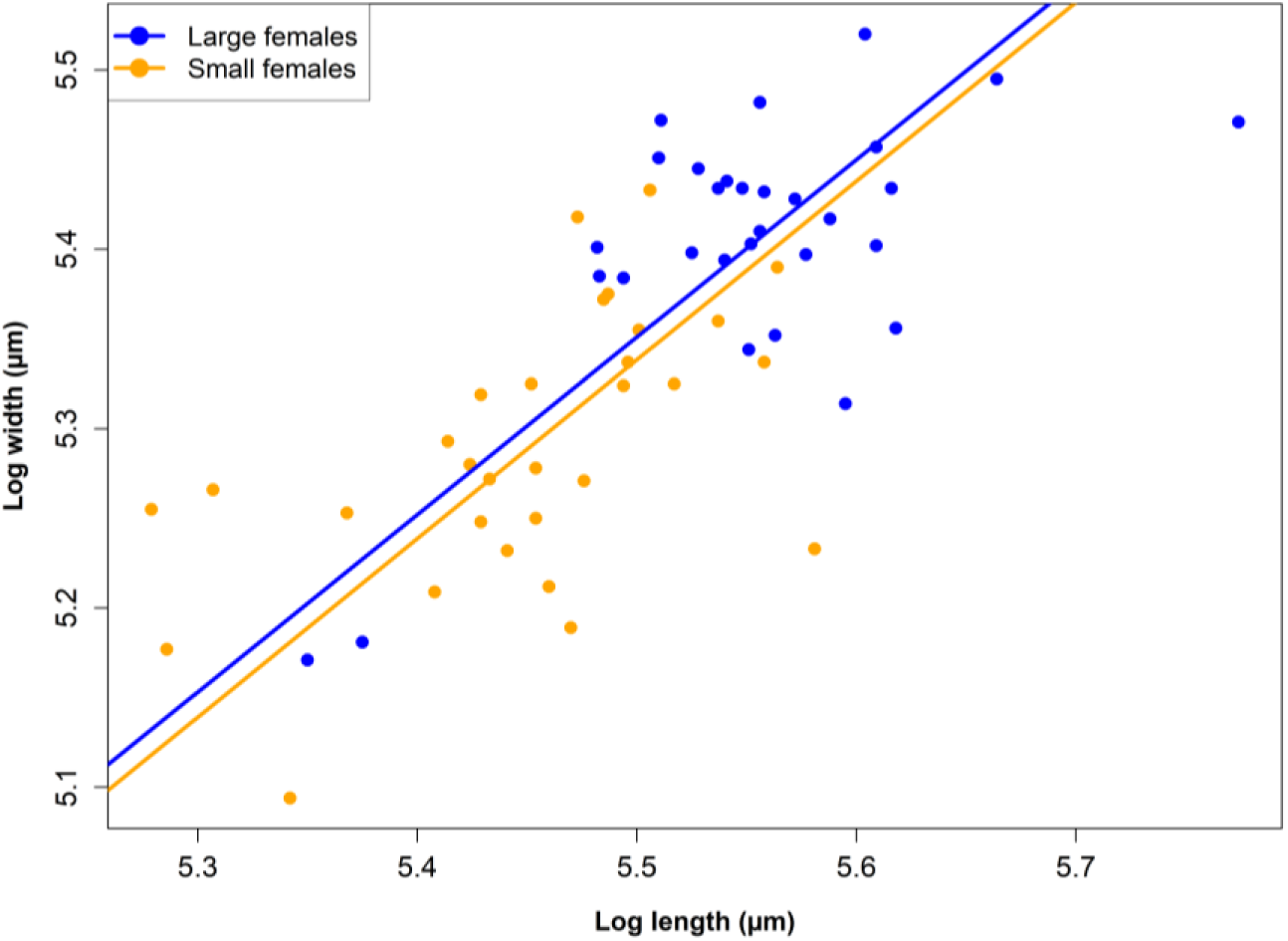
Standardized Major Axis (SMA) regression analysis representing positive trend in log (ln) length and width of categorized as "large females ≥ 4.80 mm" (*n*= 30) and "small females ≤ 4.70 mm" (*n*= 30) from antennal club of females of *I. typographus.* Each colour represents the measurements taken in length and width expressed in micrometers (µm). Blue line and dots represent the "large females" group. Orange line and dots represent the "small females" group.

### 3.3. Larger females are more responsive to (+)-isopinocamphone, whereas smaller females have higher antennal sensitivity to 1,8-cineole

EAG responses to the pheromone blend (MB:cV/ 10:1) did not differ significantly between large (*n* = 10) and small (*n* = 10) females (Exact Wilcoxon Rank Sum Test, Fig. 3A). Large females showed significantly stronger responses to four higher doses of (+)-isopinocamphone (1 µg to 1000 µg; log doses 0 to 3) (V= 48, p = 0.048; Fig. 3B). Conversely, small females exhibited significantly stronger EAG responses to three higher doses of 1,8-cineole (10 µg, 100 µg, 1000 µg; log doses 1 to 3) compared to large females (V= 7, p = 0.037; Fig. 3C). Additional details are provided in supplementary material (Table 1D).

**Figure 3.**
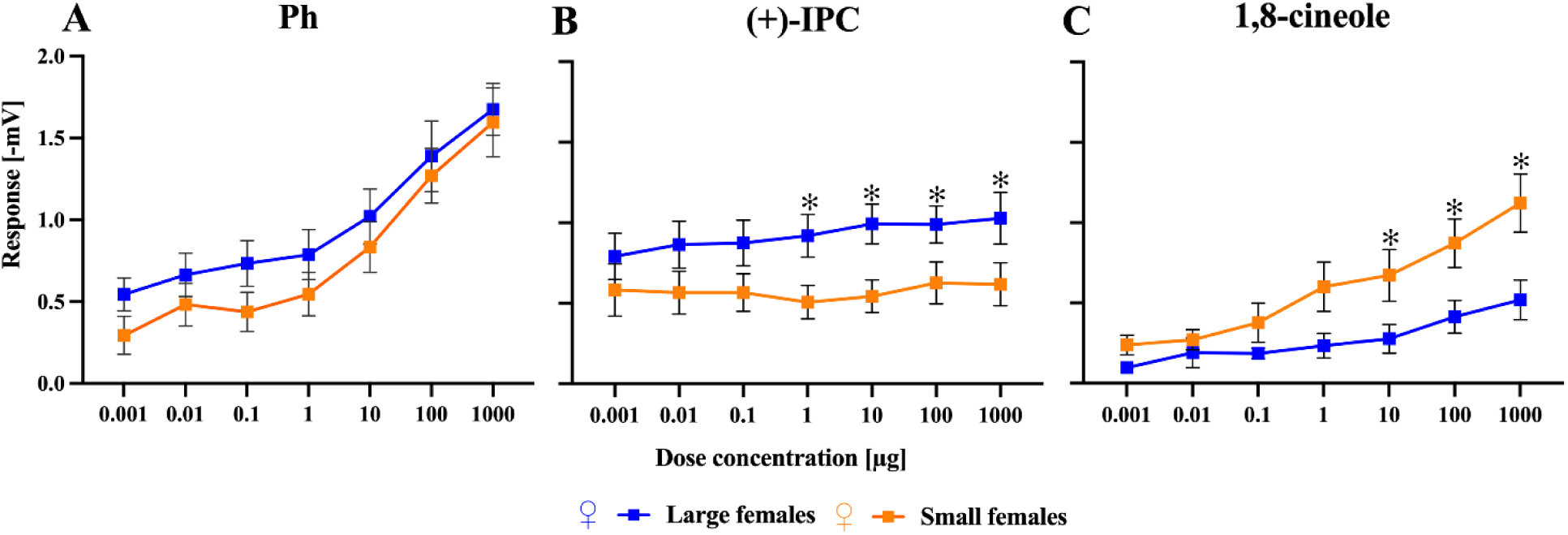
Dose–response curves based on electroantennographic (EAG) responses in female *Ips typographus*, categorized by body size: "large" (≥ 4.80 mm, *n* = 10) and "small" (≤ 4.70 mm, *n* = 10). Responses are shown for (A) Pheromone, (B) 1,8-cineole, and (C) (+)-isopinocamphone. Each compound was tested across a concentration range of 0.001 µg to 1000 µg, with hexane serving as the solvent control. EAG responses are expressed as the mean amplitude of antennal depolarizations (in millivolts), normalized by subtracting the response to hexane (blank). Error bars represent the standard error of the mean (SEM). Asterisks (*) indicate statistically significant differences between size groups at individual doses, based on the Exact Wilcoxon Rank Sum Test (*p* < 0.05).

## 4. Discussion

Our findings demonstrate that large and small female *I. typographus* respond differently to two ecologically relevant oxygenated spruce monoterpenes, 1,8-cineole and (+)-isopinocamphone, which serve as a pheromone inhibitor and a pheromone synergist, respectively. These semiochemicals influence female attraction and decision-making, with clear size-dependent variation in both antennal responses and field behavior.

### 4.1. Enhanced sensitivity and attraction of larger *Ips typographus* females to (+)-isopinocamphone may facilitate the selection of higher-quality host trees

In our field experiments, larger females were significantly more attracted to high doses of (+)- isopinocamphone, a compound known to synergize pheromone attraction, when it was presented alongside the aggregation pheromone. This behavioral pattern was supported by electroantennography (EAG) analyses, which showed that larger females exhibited stronger olfactory responses to high doses of (+)-isopinocamphone compared to smaller females. Morphometric analysis further revealed that larger females possess proportionally broader and longer antennal clubs. This increased antennal surface area likely improves their odour detection capabilities. Across various insect taxa, a correlation between antennal size and odour sensitivity has been widely documented (Makarova et al. 2022; Spaethe et al. 2007; Elgar et al. 2018; Lockey and Willis 2015). Longer antennae can house longer sensilla with greater pore density, which enhances the detection of odorants and the resulting neural activation (Mohebbi et al. 2022; Steinbrecht 2007; Liu et al. 2021). Miniaturized insects often display reductions in the antennomere number and sensilla count, as well as have shorter sensilla (Makarova et al. 2022; Steinbrecht 2007), although the diversity of sensilla types is typically maintained, allowing the detection of ecologically relevant odors (Polilov 2015; Diakova and Polilov 2020). These structural traits likely contribute to the higher sensitivity to (+)-isopinocamphone observed in larger females in our study.

The ecological implications of stronger attraction to (+)-isopinocamphone in larger females are somewhat speculative but may confer adaptive advantages. Notably, several symbiotic ophiostomatoid fungi associated with bark beetles, *G. penicillata*, *L. europhioides*, and *O. bicolor* can metabolize host tree monoterpenes into substantial quantities of (+)- isopinocamphone (Kandasamy et al. 2023). An increased olfactory response to this compound could help larger, dominant females locate trees where fungal symbionts have already detoxified monoterpenes, thereby increasing the likelihood of successful colonization. This relationship may also benefit the fungi. Larger females tend to excavate longer galleries and transport greater fungal spore loads, potentially enhancing both fungal dispersal and establishment (Foelker and Hofstetter 2014; Sallé and Raffa 2007; Sallé et al. 2005). These dynamics suggest a potential feedback loop in which fungal metabolites selectively attract the most fecund or competitive beetles, reinforcing mutualistic interactions. Future research should investigate whether larger *I. typographus* females exhibit specific preferences for fungal species producing (+)-isopinocamphone and how this might shape the evolution of beetle–fungus mutualisms. Additionally, natural enemies of *I. typographus* are responsive to isopinocamphone (Pettersson and Boland 2003), suggesting potential, yet unexplored, tri-trophic interactions linking beetle body size, fungal volatiles, and predator attraction (Souza et al. 2024; Wegensteiner et al. 2015). Trap data also indicated a higher proportion of large females in 2022 compared to 2019 caught to treatments, even the mean size of all females caught to pheromone-only was the same in both years. We attribute a shift in the proportion of large females caught to treatments to the transition from the endemic bark beetle population in 2019 to the epidemic population that occurred in 2022. During endemic periods, selective pressure on host quality increases, potentially favouring larger females that can better discriminate among semiochemical cues (Sallé et al. 2005). These findings suggest that total body length-linked behavioral strategies are modulated by population density.

### 4.2. The unexpectedly heightened antennal sensitivity of smaller ***Ips typographus*** females to 1,8-cineole contrasts with their higher behavioral attraction to this anti-attractant

An interesting contradiction was observed in the response of smaller females to 1,8-cineole. Although this compound is well-documented as an anti-attractant and toxic to *I. typographus* (Andersson et al. 2010; Jirošová et al. 2022; Zaman et al. 2024), and its addition significantly reduced the overall number of beetles captured in our field experiment to pheromone (Moliterno et al. 2024), a size-dependent pattern in females was observed. Specifically, we observed a higher proportion of smaller females in catches when exposed to high doses of 1,8-cineole combined with pheromone, compared to larger females. This pattern could still align with the antennal size hypothesis discussed earlier: those larger females with larger antennae, may detect the anti-attractant more effectively and are, therefore, more strongly repelled. However, contrary to expectations, electroantennographic (EAG) data showed that smaller females exhibited greater antennal sensitivity to 1,8-cineole than larger females despite having shorter and narrower antennal clubs.

One possible explanation for increased antennal sensitivity is that cineole-sensitive olfactory sensory neurons (OSNs) are co-localized with pheromone (*cis*-verbenol)-sensitive neurons, which suppress pheromone detection at high doses of 1,8-cineole (Andersson et al. 2010). Another explanation is that, while differences in peripheral sensitivity at the antennal surface between small and large females are influenced by antennal morphology, their host choice and decision-making may be shaped by higher-order processing in CNS regions, such as the mushroom bodies and lateral horns. These central brain areas integrate olfactory input with learning, memory, and behavioral context. Insects’ age, mating status, and energy reserves can influence both odor detection and the downstream processing of olfactory signals (Wiesel et al. 2022, Anton et al. 2007; Bodin et al. 2008; Martin et al. 2011). Consequently, the same odor may trigger different or even opposite behaviors within the same species, depending on the individual’s internal factors. Smaller females may have stronger neural connections mediating responses between cineole-sensitive OSNs and central brain regions, such as the lateral horn, which controls avoidance behaviors and suppresses their repulsive responses.

A potential ecological rationale for this pattern is linked to findings that trees with higher levels of 1,8-cineole are generally more resistant to bark beetle attacks (Schiebe et al. 2012). Larger *I. typographus* females, typically associated with higher fitness and greater capacity to kill trees (Grodzki 2004), may actively avoid such trees, recognizing them as poor-quality hosts. Doing so may increase their chances of successful colonization and reproduction (Raffa et al. 2016). In contrast, smaller females, less competitive during outbreaks, may tolerate trees with high 1,8-cineole levels as a form of competitive escape. This strategy allows them to occupy less suitable trees while avoiding competition from larger females despite the higher risks posed by the compound’s toxicity. Since our experiments were conducted in June-July, when the nutritional feeding of beetles and sister brood females from the first generation may overlap with the emergence of second-generation beetles searching for new hosts, we cannot precisely narrow down the ecological explanation solely to females seeking mates alongside suitable trees. However, we expect that the ecological principle of finding suitable host trees, where large females prefer trees with compromised defense, while smaller females avoid competition, will also apply to secondary-emerging and sister-brooding females.

### 4.3. Expanding future research framework to include males

Our analysis focused exclusively on females due to their central role in reproduction and host colonization. Additionally, field captures showed a female-biased sex ratio in traps baited with either synthetic oxygenated monoterpenes combined with pheromone or pheromone alone. This pattern is consistent with earlier reports of female-biased attraction to both aggregation pheromone (Franklin et al. 2000; Schlyter et al. 1987) and 1,8-cineole (Jirošová et al. 2022). However, it is also important to consider the potential implications for males. As the pioneer sex, males initiate host colonization and benefit from detecting semiochemicals related to the host tree’s nutritional quality and the tree’s defense ability. Unfortunately, in our catches of beetles with (+)-isopinocamphone and 1,8-cineole, there weren’t enough males for body length measurements after sorting by sex through dissection, making it impossible to obtain a statistically significant dataset. However, similarly to the total beetle catches, we observed that more males were attracted to the combination of (+)-isopinocamphone and pheromone (Moliterno et al., 2023) than to pheromone alone and in contrast, fewer males were attracted to the mixture of cineole and pheromone compared to pheromone alone.

Interestingly, when focusing on the sex ratio, we found a lower proportion of males attracted to (+)-isopinocamphone than to 1,8-cineole. However, previous studies have identified olfactory sensory neuron classes in males and females of *Ips typographus* that are primarily tuned to both (+)-isopinocamphone and 1,8-cineole (Andersson et al., 2009; Kandasamy et al., 2023), suggesting that males are equally sensitive on the periphery to both compounds as females. To better understand the signal processing in the beetle’s olfactory system and the ecological relevance of our findings, further research on sex-specific electrophysiological responses, as well as size-dependent behavior and detection abilities in males, is needed.

## 5. Conclusion

Our study demonstrates how body size influences adaptive responses in semiochemical-mediated host selection among female bark beetles. We report clear size-dependent olfactory and behavioral strategies in female *I. typographus*, linking antennal morphology, olfactory sensitivity, and host-selection behavior. The de novo spruce-derived oxygenated monoterpene 1,8-cineole and the multisource-derived hydroxylated (+)-isopinocamphone, with their contrasting ecological roles as pheromone synergist or inhibitor, respectively, may significantly influence responses in *I. typographus* females based on their size. Larger females exhibited greater olfactory sensitivity and attraction to (+)-isopinocamphone, allowing them to more effectively discriminate between suitable, more stressed, and/or fungus-colonized hosts. In contrast, smaller females were less repelled and, surprisingly, more antennally sensitive to 1,8-cineole, possibly reflecting an alternative strategy to avoid competition with larger females by exploiting lower-quality or riskier habitats. These findings suggest that body size can influence olfactory detection and subsequent CNS processing, leading to behavioral decision-making that may impact reproductive success and population dynamics of bark beetles. While our focus was on females due to their crucial role in reproduction and colonization, future research should investigate whether similar size-dependent responses occur in males.

## Authors’ contributions

AACM conceived and designed the study experiments. JS assisted in insect maintenance and rearing. AACM and MKS performed morphometric and electrophysiological analyses and collected data. AACM performed modeling work and statistical analysis of output data. SB prepared the figures and tables. AACM wrote the first draft. AACM, MKS, SB, and JS edited the draft. AJ and JH supervised the work, edited text, and provided valuable feedback.

## Acknowledgements

We are grateful to Prof. Rikard Unelius, Linnaeus university, Sweden for providing (+)- isopinocamphone for experiments. We thank Katerina Beránková for the field data from 2019. We also would like to thank both doctoral students, Rajarajan Ramakrishnan and Strádal

Jaroslav for their assistance during the sex determination of beetles, and Jaromir Bláha for managing beetle infested logs.

## Ethics approval (include appropriate approvals or waivers)

Ethical approval was not required for this study. We have performed all beetle experiments that comply with the ARRIVE guidelines and were carried out in accordance with (Scientific Procedures) Act, 1986 and associated guidelines, EU Directive 2010/63/EU for animal experiments.

## Consent to participate (include appropriate statements)

Not applicable

## Consent for publication (include appropriate statements)

All authors gave their informed consent to this publication and its content.

## Supplementary data

Supplementary data are provided as an aditional file.

