## Supplementary material for "Size-dependent behavioral and antennal responses to doses of (+)-isopinocamphone and 1,8-cineole mixed with pheromone: a potential host selection strategy in female *Ips typographus* L.": Suplementary figure and tables

**Supplementary Figure 1A:Relative catches and sex ratio of *Ips typographus* in two-year experiment of pheromone traps (alone) and baited with oxygenated host tree compounds 1,8-cineole and (+) isopinocamphone in 2019 and 2022**

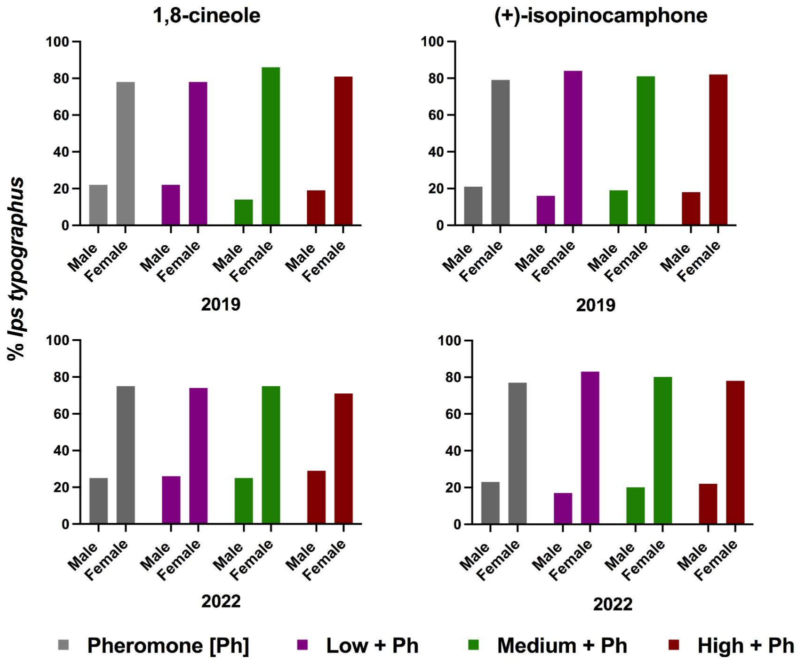


**Supplementary table 1A: Sex-ratio *I. typographus* captured in response to three doses (low, medium, high) of 1,8-cineole and (+)-isopinocamphone, along with a control (pheromone only = Ph), in 2019 and 2022. Colors represent different doses or the control**

| **1,8-cineole 2019** | **Males (%)** | **Females (%)** | total (absolute catches) |
|---|---|---|---|
| Pheromone [Ph] | 22 | 78 | 5248 |
| Low + Ph | 19 | 81 | 2333 |
| Medium + Ph | 14 | 86 | 3415 |
| High + Ph | 22 | 78 | 852 |

| **+)-isopinocamphone 201** | **Males (%)** | **Females (%)** | total (absolute catches) |
|---|---|---|---|
| Pheromone [Ph] | 21 | 79 | 5248 |
| Low + Ph | 16 | 84 | 4648 |
| Medium + Ph | 19 | 81 | 4732 |
| High + Ph | 18 | 82 | 6353 |

| **1,8-cineole 2022** | **Males (%)** | **Females (%)** | total (absolute catches) |
|---|---|---|---|
| Pheromone [Ph] | 26 | 74 | 1983 |
| Low + Ph | 25 | 75 | 1330 |
| Medium + Ph | 29 | 71 | 1269 |
| High + Ph | 25 | 75 | 263 |

| **+)-isopinocamphone 202** | **Males (%)** | **Females (%)** | total (absolute catches) |
|---|---|---|---|
| Pheromone [Ph] | 17 | 83 | 1572 |
| Low + Ph | 20 | 80 | 1267 |
| Medium + Ph | 22 | 78 | 1406 |
| High + Ph | 23 | 77 | 1917 |

**Supplementary Material, table 1B. Female of Ips typographus body length in millimeters trapped in pheromone traps in combination with 1,8-cineole or (+)-isopinocamphone in 2019 and 2022 (Anova followed by Tukey's HSD test (p = 0.05). Low (L), Medium (M) and High (H) represent relative dose trapped in combination with pheromone (Ph) versus Ph alone.**

| Anova | Df | Sum Sq | Mean Sq | F value | Pr(>F) | |
|---|---|---|---|---|---|---|
| treatments (1,8-cineole 2019) | 3 | 8.7500 | 2.9160 | 3 | 0.026 | * |
| Residuals | 196 | 182 | 0.9260 | | | |

| Anova | Df | Sum Sq | Mean Sq | F value | Pr(>F) | |
|---|---|---|---|---|---|---|
| treatments (1,8-cineole 2022) | 3 | 26 | 8.695 | 10 | <0.0001 | *** |
| Residuals | 196 | 177.650 | 0.906 | | | |

| Doses: 1,8-cineole (2019) | Estimate | Standard error | *t* value | Pr (>\|*t*\|) | |
|---|---|---|---|---|---|
| Ph + H **x** Ph | -0.5229 | 0.1920 | -2.7170 | **0.004** | * |
| Ph + M **x** Ph | -0.0223 | 0.1920 | -0.1160 | 0.999 | |
| Ph + L **x** Ph | -0.1949 | 0.1920 | -1.0130 | 0.742 | |
| Ph + M **x** Ph + H | 0.5007 | 0.1920 | 2.6010 | **0.048** | * |
| Ph + L **x** Ph + H | 0.3280 | 0.1920 | 1.7040 | 0.354 | |
| Ph + L **x** Ph + M | -0.1726 | 0.1920 | -0.8970 | 0.808 | |

| Doses: 1,8-cineole (2022) | | | | | |
|---|---|---|---|---|---|
| Ph + H **x** Ph | -0.875 | 0.1900 | -4.600 | **< 0.0001** | *** |
| Ph + M **x** Ph | -0.856 | 0.1900 | -4.499 | **< 0.0001** | *** |
| Ph + L **x** Ph | -0.746 | 0.1900 | -3.918 | **0.0007** | *** |
| Ph + M **x** Ph + H | 0.019 | 0.1900 | 0.101 | 0.999 | |
| Ph + L **x** Ph + H | 0.129 | 0.1900 | 0.682 | 0.903 | |
| Ph + L **x** Ph + M | 0.110 | 0.1900 | 0.581 | 0.937 | |

| Anova | Df | Sum Sq | Mean Sq | F value | Pr(>F) | |
|---|---|---|---|---|---|---|
| treatments ((+)-isopinocamphone 2019) | 3 | 8.2500 | 2.7480 | 3 | **0.0320** | * |
| Residuals | 196 | 177.29 | 0.1 | | | |

| Anova | Df | Sum Sq | Mean Sq | F value | Pr(>F) | |
|---|---|---|---|---|---|---|
| treatments ((+)-isopinocamphone 2022) | 3 | 6.7200 | 2.24 | 3 | **0.033** | * |
| Residuals | 196 | 148.82 | 0.7590 | | | |

| Doses: (+)-isopinocamphone (2019) | Estimate | Standard error | *t* value | Pr (>\|*t*\|) | |
|---|---|---|---|---|---|
| Ph + H **x** Ph | 0.064 | 0.0449 | 1.424 | 0.486 | |
| Ph + M **x** Ph | 0.006 | 0.0449 | 0.134 | 0.999 | |
| Ph + L **x** Ph | -0.070 | 0.0449 | -1.558 | 0.405 | |
| Ph + M **x** Ph + H | -0.058 | 0.0449 | -1.291 | 0.570 | |
| Ph + L **x** Ph + H | -0.134 | 0.0449 | -2.982 | **0.017** | * |
| Ph + L **x** Ph + M | -0.076 | 0.0449 | -1.517 | 0.331 | |

| Doses: (+)-isopinocamphone (2022) | | | | | |
|---|---|---|---|---|---|
| Ph + H **x** Ph | 0.032 | 0.04147 | 0.772 | 0.867 | |
| Ph + M **x** Ph | 0.004 | 0.04147 | 0.096 | 1.000 | |
| Ph + L **x** Ph | -0.086 | 0.04147 | -2.074 | 0.165 | |
| Ph + M **x** Ph + H | -0.028 | 0.04147 | -0.675 | 0.906 | |
| Ph + L **x** Ph + H | -0.118 | 0.04147 | -2.845 | **0.025** | * |
| Ph + L **x** Ph + M | -0.09 | 0.04147 | -2.17 | 0.135 | |

**Legend**

| | |
|---|---|
| ***Ph =*** | Pheromone |
| ***Ph + L =*** | Low released ratio |
| ***Ph + M =*** | Medium released ratio |
| ***Ph + H =*** | High released ratio |

**Supplementary Material, table 1C. Standardized major axis (SMA) regression between antennal club length and width log-transformed.**

**In bold indicating slopes close to 1 (Large: 0.99, Small: 1.0), isometrical development. R-squared showing the correlation between length and width indicated significant and positive correlations ($p \leq 0.001$) and (Small= $R^2$= 0.32, p= 0.001)**

| SMA_Group_antennal club_size | Parameter | Estimate | Lower Limit | Upper Limit | P-value |
|---|---|---|---|---|---|
| Large_females | elevation | -0.09334208 | -1.70663684 | 1.51995268 | |
| | *slope* | **0.99** | 0.741138 | 1.3221391 | |
| | R-squared | 0.43 | | | **0.0001** |
| Small_females | elevation | -0.1440909 | -1.8781666 | 1.5899849 | |
| | *slope* | **1.00** | 0.7282489 | 1.3644331 | |
| | R-squared | 0.32 | | | **0.001** |

**Supplementary Material, table 1D: Dose responses from pheromone (MB:cV; 10:1), 1,8-cineole and (+)-isopinocamphone via EAG analysis of females of *Ips typographus* categorized by body length as large (N=10) and small (N=10) obtained from F0 generation (wild generation).**
**In bold significant differences are expressed via Wilcoxon signed rank test at p = 0.05.**

| **Dose responses** *Pheromone* (MB:cV; 10:1) | **Female size** | **Average** (Standard deviation) | **Exact Wilcoxon rank sum test** | **N= Small** | **N= Large** |
|---|---|---|---|---|---|
| *1ng* | Large | 0.544 (SD= 0.32) | V=45, p = 0.083 | | |
| | Small | 0.294 (SD= 0.37) | | | |
| *10ng* | Large | 0.664 (SD= 0.42) | V=38, p = 0.322 | | |
| | Small | 0.483 (SD= 0.41) | | | |
| *100ng* | Large | 0.734 (SD= 0.44) | V=38, p = 0.322 | | |
| | Small | 0.438 (SD= 0.38) | | | |
| *1µg* | Large | 0.788 (SD= 0.49) | V=34, p = 0.556 | 10 | 10 |
| | Small | 0.546 (SD= 0.42) | | | |
| *10µg* | Large | 1.021 (SD= 0.53) | V=37, p = 0.375 | | |
| | Small | 0.834 (SD= 0.49) | | | |
| *100µg* | Large | 1.389 (SD=0.68) | V=33, p = 0.625 | | |
| | Small | 1.270 (SD=0.53) | | | |
| *1mg* | Large | 1.675 (SD= 0.50) | V=32, p = 0.695 | | |
| | Small | 1.597 (SD= 0.66) | | | |

| **Dose responses** *1,8-cineole* | **Female size** | **Average** (Standard deviation) | **Exact Wilcoxon rank sum test** | **N= Small** | **N= Large** |
|---|---|---|---|---|---|
| *1ng* | Large | 0.139 (SD= 0.13) | V=16, p = 0.275 | | |
| | Small | 0.239 (SD= 0.19) | | | |
| *10ng* | Large | 0.337 (SD= 0.53) | V=27, p = 1 | | |
| | Small | 0.271 (SD= 0.19) | | | |
| *100ng* | Large | 0.139 (SD= 0.13) | V=30, p = 0.845 | | |
| | Small | 0.239 (SD= 0.19) | | | |
| *1µg* | Large | 0.234 (SD= 0.24) | V=20, p = 0.492 | 10 | 10 |
| | Small | 0.601 (SD= 0.48) | | | |
| *10µg* | Large | 0.276 (SD= 0.28) | **V=8, p = 0.048** | | |
| | Small | 0.672 (SD= 0.50) | | | |
| *100µg* | Large | 0.414 (SD=0.32) | **V=7, p = 0.037** | | |
| | Small | 0.871 (SD=0.47) | | | |
| *1mg* | Large | 0.519 (SD= 0.74) | **V=7, p = 0.037** | | |
| | Small | 1.122 (SD= 0.49) | | | |

| **Dose responses** *(+)-isopinocamphone* | **Female size** | **Average** (Standard deviation) | **Exact Wilcoxon rank sum test** | **N= Small** | **N= Large** |
|---|---|---|---|---|---|
| *1ng* | Large | 0.791 (SD= 0.45) | V = 37, p = 0.375 | | |
| | Small | 0.581 (SD= 0.51) | | | |
| *10ng* | Large | 0.863 (SD= 0.46) | V = 40, p = 0.232 | | |
| | Small | 0.565 (SD= 0.42) | | | |
| *100ng* | Large | 0.874 (SD= 0.45) | V = 41, p = 0.193 | | |
| | Small | 0.565 (SD= 0.37) | | | |
| *1µg* | Large | 0.918 (SD= 0.41) | **V = 47, p = 0.048** | 10 | 10 |
| | Small | 0.506 (SD= 0.32) | | | |
| *10µg* | Large | 0.992 (SD= 0.38) | **V = 48, p = 0.037** | | |
| | Small | 0.543 (SD= 0.32) | | | |
| *100µg* | Large | 0.989 (SD=0.36) | **V = 47, p = 0.048** | | |
| | Small | 0.626 (SD=0.41) | | | |
| *1mg* | Large | 1.027 (SD=0.50) | **V= 47, p = 0.048** | | |
| | Small | 0.618 (SD=0.42) | | | |

**Supplementary table 1E :Sex-specific catch proportions of Ips typographus across treatments involving two compounds—(+)-isopinocamphone (IPC) and 1,8-cineole (Cine)—at three dose levels (Low, Medium, High) during two field seasons (2019 and 2022).**

**Each group includes total captures (Male, Female, Total) and the Observed Male (%) proportion in each treatment.**

| Group | Male | Female | Total | Observed Male (%) | z | p-value | Conclusion |
|---|---|---|---|---|---|---|---|
| IPC.2019.Low | 80 | 420 | 500 | 16 | 15.20 | <0.001 | Significant |
| IPC.2019.Medium | 95 | 405 | 500 | 19 | 13.86 | <0.001 | Significant |
| IPC.2019.High | 90 | 410 | 500 | 18 | 14.31 | <0.001 | Significant |
| Cine.2019.Low | 79 | 421 | 500 | 16 | 15.29 | <0.001 | Significant |
| Cine.2019.Medium | 100 | 400 | 500 | 20 | 13.41 | <0.001 | Significant |
| Cine.2019.High | 76 | 301 | 500 | 21 | 11.58 | <0.001 | Significant |
| IPC.2022.Low | 85 | 415 | 500 | 17 | 14.75 | <0.001 | Significant |
| IPC.2022.Medium | 100 | 400 | 500 | 20 | 13.41 | <0.001 | Significant |
| IPC.2022.High | 110 | 390 | 500 | 22 | 12.52 | <0.001 | Significant |
| Cine.2022.Low | 130 | 370 | 500 | 26 | 10.73 | <0.001 | Significant |
| Cine.2022.Medium | 125 | 375 | 500 | 25 | 11.18 | <0.001 | Significant |
| Cine.2022.High | 75 | 183 | 258 | 29 | 6.72 | <0.001 | Significant |

In both years, for both compounds, more females were captured compared to females , and is statistically significant.

A **z-test for proportions** was conducted to determine whether the proportion of males deviated significantly from an expected equal sex ratio (50:50).

**z** = test statistic; **p-value** = significance level; significance threshold set at $p < 0.05$.

All results showed statistically significant deviations from equal proportions.

**Supplementary table 1F. Total number of male and female *Ips typographus* caught over two field seasons in response to the compounds (+)-isopinocamphone and 1,8-cineole.**

**Data are shown separately for each compound and sex. The values represent pooled beetle catches from all trap rotations (n = 7 for 2019 and n= 8 for 2022).**

| Level | Sex ratio (formula = male/(male+female) | Male% | Female% | Total (june-july) | Male (absolute catches data) | Female (absolute catches data) |
|---|---|---|---|---|---|---|
| IPC.2019.Low | 0.16 | 16 | 84 | 1508 | 241 | 1267 |
| IPC.2019.Medium | 0.19 | 19 | 81 | 2875 | 546 | 2329 |
| IPC.2019.High | 0.18 | 18 | 82 | 2477 | 446 | 2031 |
| 2019.Ph | 0.21 | 21 | 79 | 3053 | 641 | 2412 |
| Cine.2019.Low | 0.19 | 19 | 81 | 1059 | 167 | 892 |
| Cine.2019.Medium | 0.14 | 20 | 86 | 2103 | 421 | 1682 |
| Cine.2019.High | 0.22 | 22 | 78 | 377 | 76 | 294 |
| 2019.Ph | 0.22 | 22 | 78 | 3647 | 540 | 2845 |
| IPC.2022.Low | 0.17 | 17 | 83 | 1267 | 215 | 1052 |
| IPC.2022.Medium | 0.20 | 20 | 80 | 1406 | 281 | 1125 |
| IPC.2022.High | 0.22 | 22 | 78 | 1917 | 422 | 1495 |
| 2022.Ph | 0.23 | 23 | 77 | 1572 | 362 | 1210 |
| Cine.2022.Low | 0.26 | 26 | 74 | 651 | 169 | 482 |
| Cine.2022.Medium | 0.25 | 25 | 75 | 519 | 130 | 389 |
| Cine.2022.High | 0.29 | 29 | 71 | 258 | 75 | 183 |
| 2022.Ph | 0.25 | 25 | 75 | 537 | 134 | 403 |

**Supplementary table 1G: Pearson's Chi-squared test with Yates' continuity correction comparing male and female Ips typographus catches for two compounds, (+)-isopinocamphone (IPC) and 1,8-cineole across different dose levels and years.**
**Data represents absolute beetle catches pooled from the respective number of trap rotations per year (2019: 7 rotations; 2022: 8 rotations).**

| Comparison (2019) | 2-sample test for equality of proportions with continuity correction | | | | |
|---|---|---|---|---|---|
| Females | Prop1 | Prop2 | *Chi*-squared | p-value | Significance |
| IPC_Low vs Cine_Low | 0.840 | 0.842 | 0.020 | 0.88 | **ns** |
| IPC_Medium vs Cine_Medium | 0.810 | 0.799 | 0.819 | 0.36 | **ns** |
| IPC_High vs Cine_High | 0.891 | 0.794 | 1.385 | 0.23 | **ns** |

| Comparison (2019) | 2-sample test for equality of proportions with continuity correction | | | | |
|---|---|---|---|---|---|
| Males | Prop1 | Prop2 | *Chi*-squared | p-value | |
| IPC_Low vs Cine_Low | 0.159 | 0.157 | 0.020 | 0.88 | **ns** |
| IPC_Medium vs Cine_Medium | 0.189 | 0.200 | 0.819 | 0.36 | **ns** |
| IPC_High vs Cine_High | 0.180 | 0.205 | 1.381 | 0.23 | **ns** |

| Comparison (2022) | 2-sample test for equality of proportions with continuity correction | | | | |
|---|---|---|---|---|---|
| Females | Prop1 | Prop2 | *Chi*-squared | p-value | |
| IPC_Low vs Cine_Low | 0.830 | 0.740 | 21.710 | *<0.001* | *** |
| IPC_Medium vs Cine_Medium | 0.800 | 0.749 | 5.785 | *0.016* | * |
| IPC_High vs Cine_High | 0.779 | 0.709 | 6.422 | *0.011* | * |

| Comparison (2022) | 2-sample test for equality of proportions with continuity correction | | | | |
|---|---|---|---|---|---|
| Males | Prop1 | Prop2 | *Chi*-squared | p-value | |
| IPC_Low vs Cine_Low | 0.169 | 0.259 | 21.710 | *<0.001* | *** |
| IPC_Medium vs Cine_Medium | 0.199 | 0.250 | 5.785 | *0.016* | * |
| IPC_High vs Cine_High | 0.220 | 0.290 | 1.381 | *0.011* | * |

Prop= (probability)

**IPC_Male**, **IPC_Female**, **Cine_Male**, and **Cine_Female** columns indicate total captures by sex and compound.
**Chi-sq** = Chi-squared statistic; **df** = degrees of freedom; **p-value** = significance level of difference in male vs. female catch proportic
Significance thresholds: $p < 0.05$ (*), $p < 0.001$ (**).

| Group | IPC_Male | IPC_Female | Cine_Male | 'ine_Fema | Chi-sq | df | p-value |
|---|---|---|---|---|---|---|---|
| 2019 - Low | 241 | 1267 | 167 | 892 | 0.008 | 1 | 0.9285 |
| 2019 - Medium | 546 | 2329 | 421 | 1682 | 0.7552 | 1 | 0.3848 |
| 2019 - High | 446 | 2031 | 76 | 294 | 1.2174 | 1 | 0.2699 |
| 2022 - Low | 215 | 1052 | 169 | 482 | 21.1517 | 1 | <0.001 |
| 2022 - Medium | 281 | 1125 | 130 | 389 | 5.4878 | 1 | 0.0191 |
| 2022 - High | 422 | 1495 | 75 | 183 | 6.0283 | 1 | 0.0141 |

Are the proportions of females significantly different between IPC and Cine doses (Low versus Low) and so on?

**In 2019, both treatments attracted beetles with similar sex distributions, regardless of dose.**
**In 2022, IPC females distribution were significantly higher at Low, Medium or High doses than 1,8-cineole tested doses**
